# Gestational inhalation of nanoparticles disrupts placental zone structure and induces vascular placentation in rats

**DOI:** 10.64898/2026.06.08.730946

**Authors:** Talia Seymore, Destiny McWilliams, Kevin Ozkuyumcu, Pedro Louro, Chelsea Cary, Michael Goedken, Laurie Joseph, Phoebe Stapleton

## Abstract

Airborne contaminants represent a significant environmental health concern for vulnerable populations, including pregnant individuals. In particular, maternal inhalation of particulate matter (PM) during pregnancy has been linked to adverse outcomes such as fetal growth restriction (FGR). Increasing evidence identifies placental dysfunction as a mechanism for this condition. Placental efficiency, defined as the ratio of fetal mass to placental mass, is frequently altered in FGR. Many aspects contribute to placental efficiency including surface area available for nutrient and waste exchange and placental vascularization. In this study, we hypothesized that maternal inhalation of ultrafine PM during pregnancy would reduce the size and/or number of placental structures that are necessary for nutrient transport. Engineered titanium dioxide nanoparticles (nano-TiO_2_) were used as a proxy for ultrafine PM and pregnant Sprague Dawley rats were exposed via whole-body inhalation to nano-TiO_2_ aerosols (9.23 ± 0.39 mg/m^3^) from gestational day (GD) 5 to 19. On GD 20, placentas were collected and processed for histological evaluation. While gestational inhalation of nano-TiO_2_ did not affect placental weight or efficiency, it reduced decidua and labyrinth zone size. Exposed placentas exhibited compensatory adaptations characterized by increased blood space number and maternal blood space expansion. Together, these findings indicate that inhalation of nanoparticles disrupts placental structure while simultaneously eliciting adaptive vascular responses that may preserve nutrient exchange capacity. By characterizing the effects of PM exposure on placental morphology and structure, this study highlights the placenta as a vulnerable target of inhaled pollutants and provides mechanistic insight into pathways contributing to PM-induced FGR.

## Introduction

Airborne pollutants are an environmental health concern for many sensitive populations, including pregnant people. Specifically, inhalation of aerosolized particulate matter (PM) during pregnancy is associated with adverse fetal development and pregnancy complications [1-6]. Many health concerns arising from inhaling PM stem from its effects on the respiratory and cardiovascular systems, largely due to the inflammation and oxidative stress these particles induce in biological systems [6-11]. Furthermore, ultrafine PM, characterized as being less than 100 nm in diameter (i.e., nanoparticles), are of particular concern due to their ability to cross the air-blood barrier in the lungs and translocate to and deposit within distal organs, including those of the reproductive system [12-15]. The etiology for pregnancy-related consequences remains unclear; however, a major concern is how toxicological components of PM affect the placenta. The placenta is a temporary organ that develops during pregnancy to regulate nutrient transport and waste exchange for the developing fetus, and fetal health is directly dependent on placental function [16]. Therefore, the placenta may play a critical role in PM-induced adverse fetal developmental outcomes, such as fetal growth restriction (FGR).

Placental vasculature is essential for the transport of growth-related nutrients from the maternal blood into the fetal circulation. Because the human placenta is hemochorial, maternal blood can directly bathe over the cellular barrier of the placenta called syncytiotrophoblasts. In the human placenta, these cells form villous structures that line “maternal blood spaces” and serve as the site of nutrient and waste exchange. On the fetal side of this barrier, fetal capillaries or “fetal blood spaces” that are lined by fenestrated endothelial cells will receive nutrients [17]. Taken together, the establishment of these structures dictate nutrient transport, placental efficiency, and fetal access to nutrients. Placental efficiency is defined as the grams of fetus produced per gram of placenta; therefore, placental inefficiency is commonly implicated in FGR [18]. It can be altered by changes in surface area for exchange, arrangements in fetal and maternal blood spaces, and intrahemal barrier thickness, or the thickness of the syncytiotrophoblast layer that separates the maternal and fetal blood space [19]. Thicker barriers between the maternal and fetal circulations, reduced surface area for exchange, and reduced placental vascularization are all associated with an inefficient placenta and consequently, growth restricted fetuses [19]. Several human and *in vivo* studies support an association between exposure to PM and placental dysfunction including dysregulated vascularization, impaired trophoblast invasion, alterations in placental ultrastructure, and changes to placental weight [20-23]. These studies have a specific focus on heterogeneous PM of varying particle sizes and chemical composition; therefore, it is unclear what toxicological components are driving these outcomes.

In the rodent placenta, the labyrinth zone, the area of nutrient and waste exchange also represents the fetal-facing zone. It is the largest zone in the rodent placenta and it is highly vascularized [24]. To establish this network with the maternal compartment, the extravillous trophoblast cells of the placenta migrate into the spiral arteries of the uterine endometrium [25]. This site of invasion, the decidua, and the receptivity of this space determines the success of spiral artery remodeling. Accordingly in the rodent placenta, the metrial gland and collectively with the decidua zone, these are maternal-facing structures that regulate trophoblast migration [24, 26]. Lastly, the junctional zone, present in the center of the labyrinth and decidua zone, is important for glycogen storage and hormone secretion [24].

In this study, we investigated the impact of PM on placental structure and vascularization. To isolate the direct effects of systemic PM independent of chemical composition, we utilized engineered titanium dioxide nanoparticles (nano-TiO_2_) as a model of ultrafine PM. This approach allows us to bypass outcomes that can be due to particle chemistry and focus on particle-driven disruptions in placental function. We hypothesized that maternal inhalation of nano-TiO_2_ during pregnancy would reduce the size and/or number of placental structures that are necessary for nutrient transport and waste exchange. Using a rat model, we assessed multiple compartments including the labyrinth, junctional, and decidua zones. By determining the impact of PM exposure on placental gross structure and ultrastructure, this study identifies the placenta as a vulnerable target and provides insight into mechanisms underlying PM-induced FGR. Moreover, these structural alterations provide rationale for investigating molecular adaptations that may arise in response to placental dysfunction.

## Methods

### Animals and Exposure

Twenty-four pregnant Sprague Dawley rats were obtained from Charles River Laboratories (Kingston, NY) and delivered on gestational day (GD) 4 to an AAALAC-accredited vivarium where animals had *ad libitum* access to food and water. All experimental protocols were approved by the Rutgers Institutional Animal Care and Use Committee. Following a 24-hr acclimation period, animals were randomly assigned to exposed groups (n=12) or naïve control groups (n=12) that did not enter the inhalation facility. Dams were exposed to an average concentration of 9.23±0.39 mg/m^3^ of aerosolized nano-TiO_2_ (129 ± 1.84 nm) powder (Aeroxide TiO_2_, Parsippany, NJ). Exposures were conducted 4-5 hr/d, 5d/wk from GD 5 to GD 19 using a custom-built 84 L whole-body inhalation chamber (Figure 1) (IEStechno, Morgantown, WV) as previously described [27-31]. A target exposure concentration of 10 mg/m^3^ was selected to approximate the maximum permissible exposure limits for occupational airborne chemical contaminants (Title 8, Article 107) [32].

**Figure 1.**
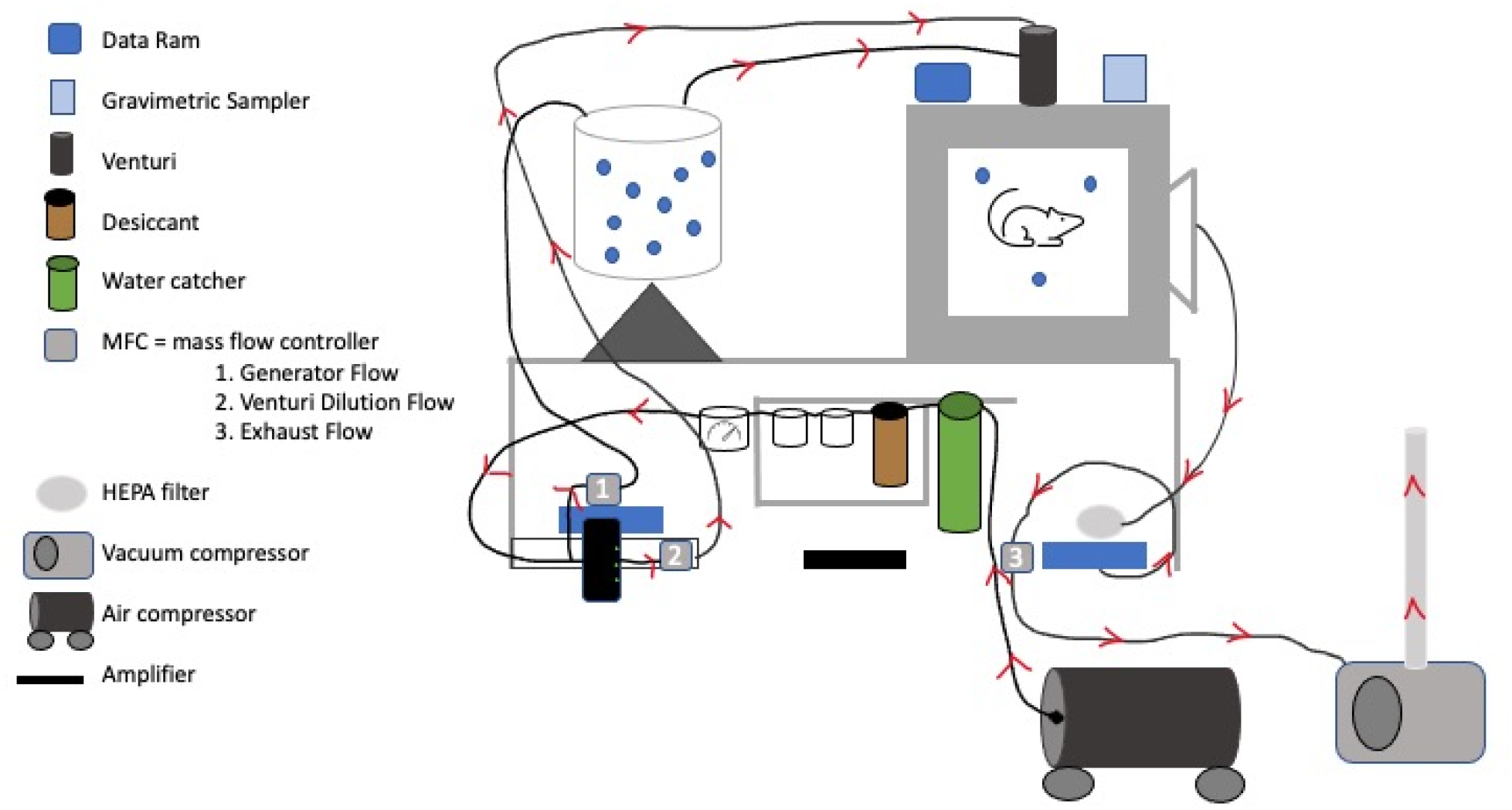
Custom-built 84 L whole-body inhalation facility. Figure created by graduate student Samantha Adams.

### Tissue Collection

On the morning of GD 20, dams were anesthetized with isoflurane (5% induction, 3% maintenance) and euthanized via pneumothorax and heart removal. Uterine horns were isolated, removed, and dissected to reveal feto-placental units. Fetal sex was determined by anogenital distance and gonad visualization. Fetal weight and placental weight were recorded and placental efficiency was calculated by dividing fetal weight by placental weight (FW/PW) (n=28). A separate group of animals were used for histological evaluation (n=12). For histological evaluation, uterine horns were opened, fetuses were removed, and sex was recorded. The remaining uterine horn with placentas attached was drop-fixed in 10% neutral buffered formalin and prepared for histological evaluation. Uterine-placental units were separated from each other and embedded in paraffin. Five µm sections were prepared and mounted on glass slides.

### Evaluation of Placental Blood Spaces and Zone Sizes

All histology samples were stained with hematoxylin and eosin (H&E) and reviewed by a third-party, board-certified veterinary pathologist for overt pathologies. H&E slides were used for the assessment of maternal and fetal blood spaces as well as labyrinth, junctional, and decidua zone sizes. Whole-slide images were acquired at 20x magnification using a Zeiss Axioscan 7 and analyzed with ZEN Blue Software. To evaluate blood space size and intrahemal barrier thickness, three maternal-fetal blood space units were identified on three sections per placenta spanning the labyrinth zone from the fetal to the maternal side. Fetal blood spaces were distinguished by endothelial lining and presence of large, immature erythrocytes, whereas maternal blood spaces contained smaller, mature erythrocytes. Blood space perimeters were traced and surface area calculated, and intrahemal barrier distance between each maternal-fetal blood space unit was recorded. Values were averaged per placenta. Blood space number was determined by acquiring three images (25000 μm^2^ each) from the labyrinth zone of each placenta. The number of maternal and fetal blood spaces in each image was recorded and averaged. For placental zone analysis, the labyrinth zone, junctional zone, and decidua zones were identified, the perimeter of each zone was traced, and surface area was calculated.

### Evaluation of Trophoblast Invasion

A subset of unstained slides was deparaffinized, rehydrated, and subjected to heat-induced antigen retrieval with citrate buffer (10 mM, pH 6.0) for 15 minutes in a steamer. Sections were blocked with 100% normal goat serum for 2 hr at room temperature. Sections were then incubated overnight at 4°C with rabbit polyclonal anti-alpha smooth muscle actin (αSMA) (ab5694, 1:1000, Abcam, Cambridge, MA). After primary antibody incubation, sections were incubated with a biotinylated goat anti-rabbit secondary antibody (BA-1000, 1:200, Vector Laboratories, Burlingame, CA) for 30 min at room temperature. αSMA smooth muscle staining was visualized using a diaminobenzidine (DAB) peroxidase substrate kit (SK-4100, Vector Laboratories, Newark, CA) and counterstained with Mayer’s hematoxylin (26043, Electron Microscopy Sciences, Hatfield, PA). Whole-slide images were acquired at 20x magnification using a Zeiss Axioscan 7 and analyzed with ZEN Blue Software. Corresponding H&E slides were used to located three blood vessels in the metrial gland of each placenta. The same blood vessels were located on αSMA-stained slides and trophoblast invasion was scored based on the percent continuity of positive αSMA expression around the vessel: score 1: 1-20% continuity, score 2: 20-40% continuity, score 3: 40-60% continuity, score 4: 60-80% continuity, and score 5: 80-100% continuity. Accordingly, less invasion is associated with a continuous layer of αSMA expression around the vessel, i.e. a score of 5. The median invasion score for each placenta was reported.

### Statistical Analyses

All data are presented as mean ± standard error mean (SEM). Statistical analyses were performed and graphs generated using GraphPad Prism v10 (Boston, MA). All data stratified by sex was analyzed by two-way ANOVA with Tukey’s post-hoc analysis. Placental weight and placental efficiency were reported as a litter average for each dam. Zone size, vessel invasion, and blood space analyses represent two placentas per dam (one male and one female).

## Results

### Placental Efficiency

Placental weight and efficiency were recorded on GD 20 for a larger cohort (n=28 litters) and reported as an overall average (Supplemental Figure 1). Exposure to nano-TiO_2_ had no effect on placental weight (Supplemental Figure 1A) nor placental efficiency (Supplemental Figure 1B).

### Placental Zone Size

Placental zone sizes were evaluated histologically. Exposure to nano-TiO_2_ significantly reduced the surface area of the labyrinth zone by 9.3% (Figure 2A), while the junctional zone was not affected by exposure (Figure 2B). In contrast, decidua zone size was significantly reduced by 16.3% in the exposed group compared to naïve controls (Figure 2C). Collectively, gestational inhalation of nano-TiO_2_ negatively impacts the primary site of maternal-fetal exchange, as well as the decidua, the maternal-facing zone.

**Figure 2.**
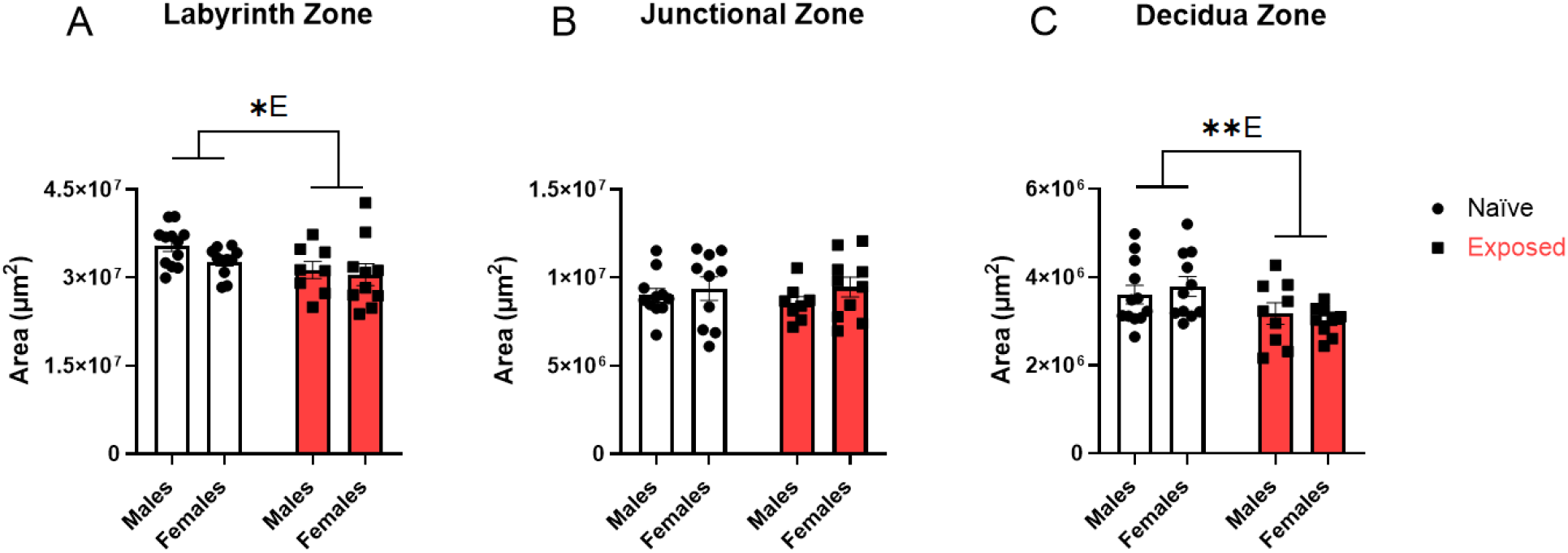
Effects of nano-TiO_2_ inhalation on placental zone surface area. The surface area of the labyrinth zone (A), junctional zone (B), and decidua zone (C) was measured histologically using ZEN Blue software on GD 20. Data are mean ± SEM, n=8-12 litters/treatment group. ^*E^Significant exposure factor (p≤0.05), **E (p<0.01); as determined by two-way ANOVA and Tukey’s post-hoc analysis.

### Placental Blood Spaces

Placental maternal and fetal blood spaces were quantified histologically (Figure 3). Exposure to nano-TiO_2_ had no effect on fetal blood space size (Figure 3A), but maternal blood space size increased by 55.7% relative to naïve controls (Figure 3B). In addition to this expansion, exposure significantly increased the number of fetal blood spaces by 36.4% (Figure 3C) and maternal blood spaces by 10.1% (Figure 3D).

**Figure 3.**
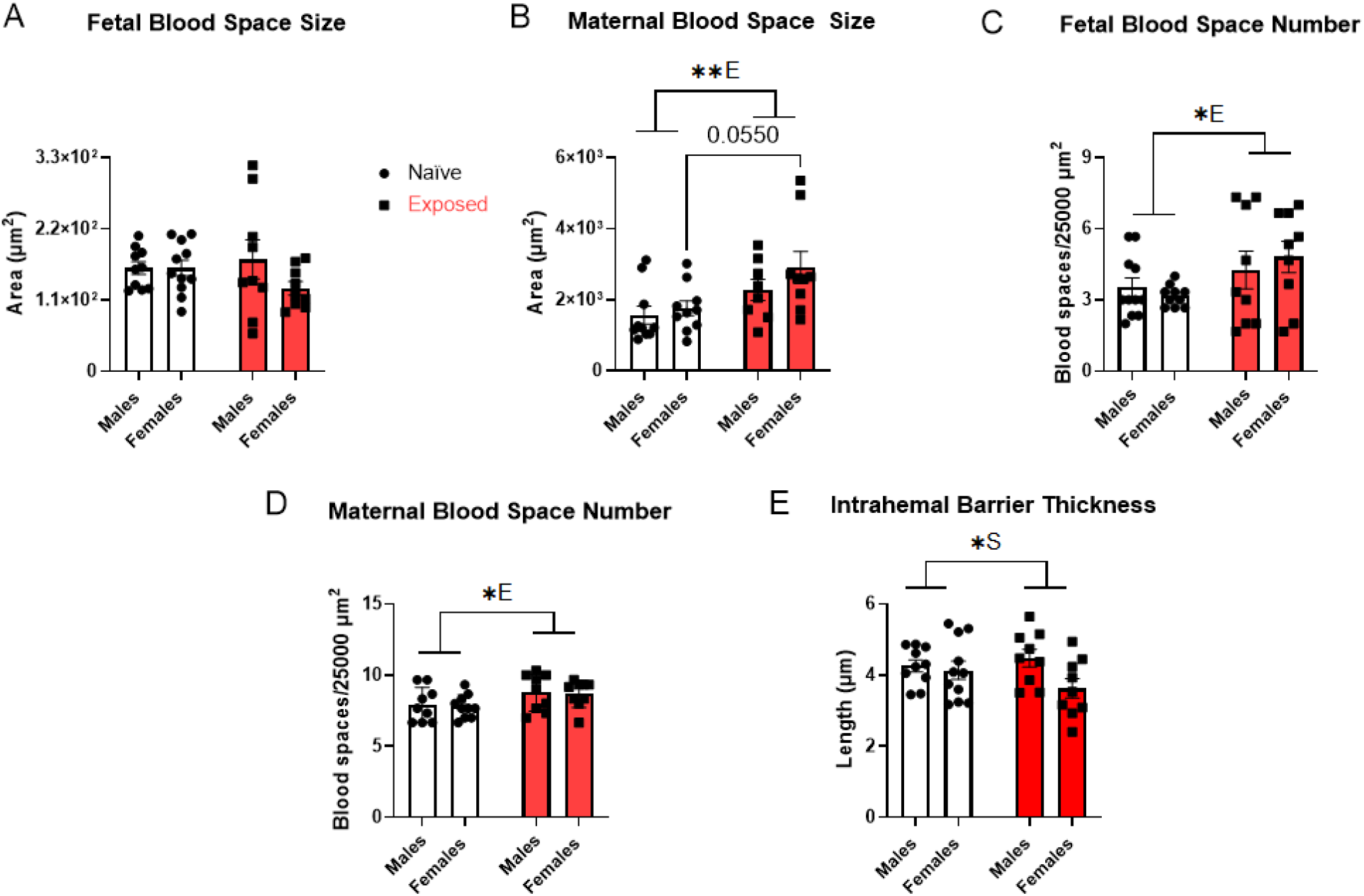
Effects of nano-TiO_2_ inhalation on placental blood spaces and intrahemal barrier thickness. Placentas were histologically evaluated for blood space size and distance between maternal-fetal blood space units on GD 20 (A-C). Blood space number was also quantified (D and E). Data was evaluated using ZEN Blue software. Data are mean ± SEM, n=9-11 litters/treatment group. ^*E^Significant exposure factor (p≤0.05), **E (p<0.01); ^*S^Significant sex factor (p≤0.05) as determined by two-way ANOVA and Tukey’s post-hoc analysis.

Intrahemal barrier thickeness was not affected by exposure; however, a sex-specific difference was observed, with male placentas exhibiting an intrahemal barrier 11.4% thicker than females (Figure 3E). Taken together, gestational inhalation of nano-TiO_2_ promoted expansion and increased density of placental blood spaces without altering intrahemal barrier thickness.

### Trophoblast Invasion

Trophoblast invasion in spiral arteries of the metrial gland was immunohistologically evaluated using an anti-αSMA antibody (Figure 4A). Exposure to nano-TiO_2_ had no significant effect on trophoblast invasion; however, exposed placentas showed a decrease in αSMA-positive cells along the spiral artery wall (Figure 4B). Accordingly, exposed placentas exhibited a trend towards a lower invasion score compared to naïve controls, suggesting slightly greater invasion of metrial gland vessels in the exposed group (Figure 4B). Interestingly, we observed a significant effect of sex, with male placentas showing a median invasion score of 2 and female placentas a median score of 3, indicating enhanced spiral artery invasion in males (Figure 2.4).

**Figure 4.**
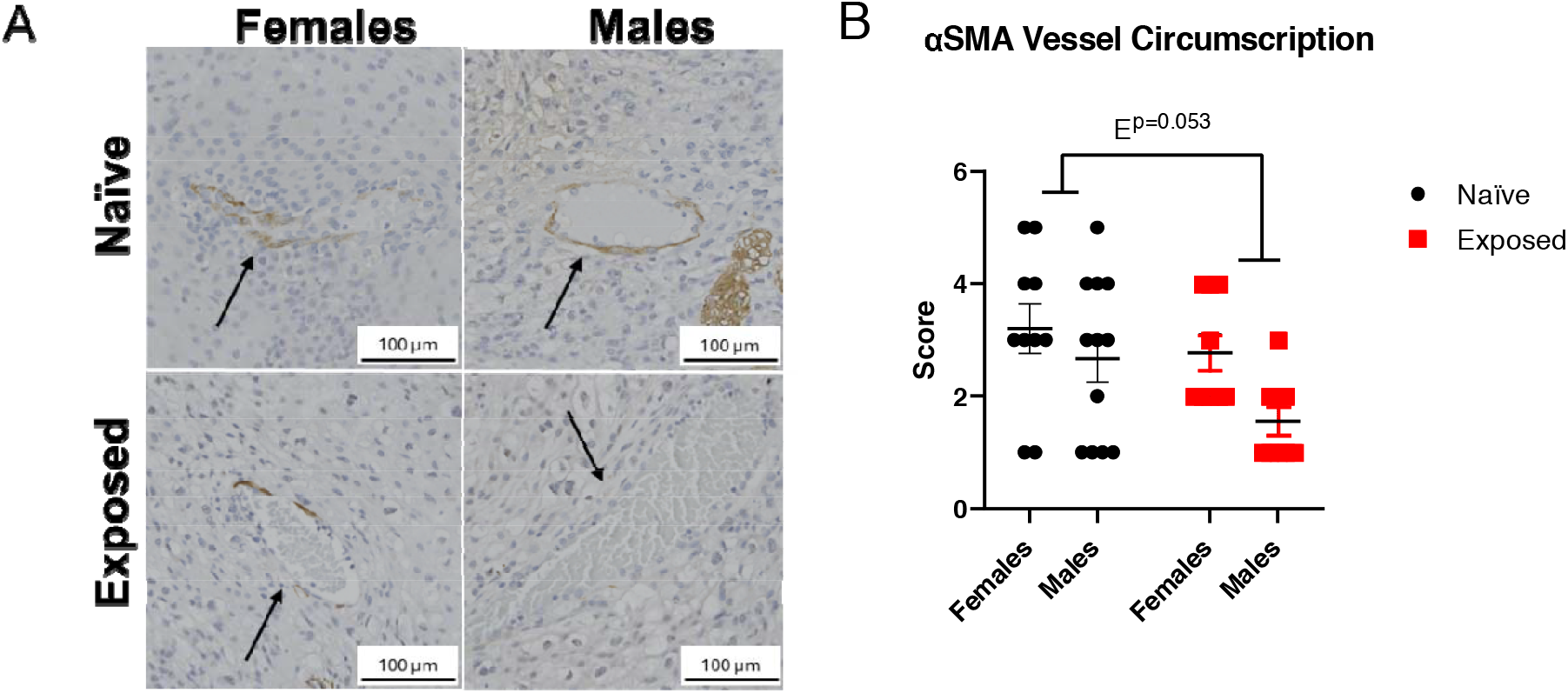
Effects of nano-TiO_2_ inhalation on trophoblast invasion. Placentas were stained with αSMA and blood vessels in the metrial gland were evaluated for positive staining as a measure of vascular invasion. Representative images were chosen based on average score of each treatment group (A). Blood vessel staining was scored on a scale of 1-5: 1 (fully invaded) to 5 (not invaded) (B). Black arrows denote blood vessels. Data are mean ± SEM, n=9-12 litters/treatment group. Analyzed by two-way ANOVA.

## Discussion

While gestational inhalation of nano-TiO_2_ did not affect placental weight or efficiency, it altered placental architecture and vascular organization. Specifically, exposure was associated with reductions in labyrinth and decidua zone size, accompanied by an increase in both maternal and fetal blood space number, as well as an expansion of maternal blood space size. Collectively, these findings suggest that the inhalation of ultrafine PM disrupts placental structural organization; however, the placenta exhibits adaptive vascular responses that may compensate for these architectural changes and rescue nutrient exchange capacity.

Coarse and fine PM exposure has been associated with reductions in placental chorionic disk area and placental weight in human studies [23, 33, 34]. Similarly, *in vivo* exposure to ultrafine PM has been linked to decreased placental weight in females, highlighting a potential sex-specific placental response to environmental stress [35]. Across many of these studies, PM exposure was also associated with an increased risk of FGR, suggesting compromised placental function and reduced placental efficiency. Although no changes in placental weight were observed in our present study, these gross measures can mask functional and molecular stress responses such as zone-specific alterations in placental architecture.

There is evidence to suggest placental responses to environmental stress are zone-specific. Specifically, the labyrinth zone is highly metabolically active and given its dense vascularization and primary role in nutrient and gas exchange, it is vulnerable to reactive oxygen species and oxidative stress [36]. In contrast, the junctional zone is primarily responsible for hormone production and may therefore be more susceptible to endocrine-disrupting insults. As the maternal-facing zone, the decidua is the first placental region to encounter circulating toxicants [24]. *In vivo* exposure to fine PM has been shown to reduce junctional zone size without affecting other placental zones [37], while exposure to ultrafine PM increased decidua zone size in a sex-specific manner [35]. In our model, gestation inhalation of ultrafine PM was associated with reductions in both labyrinth and decidua zone size. Together, these findings suggest that ultrafine PM may preferentially impact vascularized placental zones, including regions critical for spiral artery invasion. These differences may reflect particle size-dependent effects.

Adaptations in the arrangement and density of maternal and fetal vasculature are a well-established mechanism to improve placental transport capacity, especially when fetal growth is threatened [38]. In support of this, *in vivo* exposure to fine PM has been shown to increase angiogenic factors in the labyrinth zone [22] and increase fetal blood spaces [39]. Similarly, exposure to ultrafine PM also increased fetal vessel area, but only in male placentas [35]. Consistent with these adaptive responses, our data revealed increases in fetal blood vessels and expansion of maternal blood spaces in the labyrinth zone. While maternal blood spaces are not discrete vessels, their enlargement reflects structural remodeling of the maternal-fetal interface that increases surface for the maternal blood to bathe over the syncytiotrophoblast layer, supporting nutrient and waste exchange. These structural adaptations may be linked to changes in trophoblast behavior, including the regulation of spiral artery invasion.

In line with this concept, an *in vivo* study reported decreased invasion of spiral arteries in placentas exposed to fine PM [37] whereas our model using ultrafine PM exhibited a trend toward increased trophoblast invasion in the metrial gland. Although not statistically significant, it is consistent with an adaptive response aimed at enhancing vascular access and perfusion. Increased trophoblast invasion could facilitate the observed expansion of maternal blood spaces. While it is unclear what may be driving these opposing responses, there is evidence to support that hypoxic environments enhance trophoblast invasion [40, 41]. In our model, we have previously demonstrated reduced uteroplacental blood flow following ultrafine PM exposure, indicating an ischemic state that may result in reduced oxygen tension and delivery [27, 42, 43]. Together, these findings support the concept that gestational inhalation of ultrafine PM elicits placental adaptations that improve placental efficiency.

While this study provides insight into placental adaptations following exposure to PM, several limitations should be considered. First, these data lack functional measures, meaning that we cannot determine if these adaptations have beneficial consequences. Some valuable measures would include evaluating oxygen delivery and active transport of nutrients. Second, since we only evaluated toxicological effects on GD 20, we may be missing the windows of adaptation and other toxicological impacts that may occur throughout pregnancy. This is especially important for trophoblast invasion as this process begins during early placentation and peaks during mid gestation [44]. This would allow us to determine early insults as a response to ultrafine PM exposure during pregnancy. The unique physiochemical properties of nano-TiO_2_ also warrant consideration, particularly because this model cannot recapitulate the inherent heterogeneity of ambient ultrafine PM. Unlike engineered nano-TiO_2_, ambient PM carries adsorbed airborne heavy metals and organic compounds on its surface, which can likely exacerbate toxicological consequences [45].

In conclusion, the placenta is a dynamic and adaptive organ in response to environmental stress. Our findings indicate that vascular adaptations may function as a key compensatory mechanism in the face of insults that disrupt placental architecture. These results underscore the placenta as an important endpoint when evaluating the effects of gestational exposure to environmental contaminants on fetal development. These changes can have profound implications for fetal health and highlight the need for continued investigation into the placental mechanisms underlying FGR and other pregnancy complications associated with PM inhalation.

## Acknowledgments

This work was supported by the National Institutes of Health [NIH R01ES031285, T32ES007148, and P30 ES005022]. The authors also acknowledge the contributions of lab managers Tanisha Brunson-Malone and Dr. Marianne Polunas, and PhD student Samantha Adams for performing the animal exposures.

## References

1. Wilhelm, M., et al., Traffic-Related Air Toxics and Term Low Birth Weight in Los Angeles County, California. Environmental Health Perspectives, 2012. 120(1): p. 132–138.

2. Chen, M.M., et al., Influence of Environmental Tobacco Smoke and Air Pollution on Fetal Growth: A Prospective Study. Int J Environ Res Public Health, 2020. 17(15).

3. Xu, X., et al., Long-term Exposure to Ambient Fine Particulate Pollution Induces Insulin Resistance and Mitochondrial Alteration in Adipose Tissue. Toxicological Sciences, 2011. 124(1): p. 88–98.

4. Gustavsson, P., et al., Risk of preeclampsia and gestational diabetes after occupational exposure to chemicals during pregnancy-A cohort study of births in Sweden 1994-2014. Environ Res, 2025. 279(Pt 1): p. 121802.

5. Pereira, G., M.B. Bracken, and M.L. Bell, Particulate air pollution, fetal growth and gestational length: The influence of residential mobility in pregnancy. Environmental Research, 2016. 147: p. 269–274.

6. Park, S., et al., Effect of Particulate Matter 2.5 on Fetal Growth in Male and Preterm Infants through Oxidative Stress. Antioxidants, 2023. 12(11): p. 1916.

7. Kyung, S.Y. and S.H. Jeong, Particulate-Matter Related Respiratory Diseases. Tuberc Respir Dis (Seoul), 2020. 83(2): p. 116–121.

8. Li, Y., et al., Reactive oxygen species induced by personal exposure to fine particulate matter emitted from solid fuel combustion in rural Guanzhong Basin, northwestern China. Air Quality, Atmosphere & Health, 2019. 12(11): p. 1323–1333.

9. Du, Y., et al., Air particulate matter and cardiovascular disease: the epidemiological, biomedical and clinical evidence. J Thorac Dis, 2016. 8(1): p. E8–e19.

10. Bhatnagar, A., Cardiovascular Effects of Particulate Air Pollution. Annu Rev Med, 2022. 73: p. 393–406.

11. Xu, F., et al., Investigation of the chemical components of ambient fine particulate matter (PM(2.5)) associated with in vitro cellular responses to oxidative stress and inflammation. Environ Int, 2020. 136: p. 105475.

12. Cary, C.M., et al., Ingested polystyrene nanospheres translocate to placenta and fetal tissues in pregnant rats: Potential health implications. Nanomaterials, 2023. 13(4): p. 720.

13. Ragusa, A., et al., Plasticenta: First evidence of microplastics in human placenta. Environment International, 2021. 146: p. 106274.

14. D’Errico, J.N., et al., Maternal, placental, and fetal distribution of titanium after repeated titanium dioxide nanoparticle inhalation through pregnancy. Placenta, 2022. 121: p. 99–108.

15. Parikh, G., et al., Maternal exposure to PM(2.5) and its association with adverse pregnancy and birth outcomes: A prospective cohort study. Reprod Toxicol, 2025. 140: p. 109127.

16. Jarzembowski, J.A., Normal Structure and Function of the Placenta, in Pathobiology of Human Disease, L.M. McManus and R.N. Mitchell, Editors. 2014, Academic Press: San Diego. p. 2308–2321.

17. Wang, Y., Vascular Biology of the Placenta. Colloquium Series on Integrated Systems Physiology: From Molecule to Function, 2010. 2(1): p. 1–98.

18. Wilson, M. and S. Ford, Comparative aspectsof placental efficiency. Reproduction Supplement (58), 1999: p. 223–232.

19. Fowden, A.L., et al., Placental efficiency and adaptation: endocrine regulation. The Journal of physiology, 2009. 587(14): p. 3459–3472.

20. Lee, G., et al., Impact of particulate matter 2.5 on placental ultrastructure including mitochondrial damage through oxidative stress. Reprod Toxicol, 2025. 136: p. 108973.

21. Hur, H., et al., Maternal Exposure to Diesel Exhaust Particles (DEPs) During Pregnancy and Adverse Pregnancy Outcomes: Focusing on the Effect of Particulate Matter on Trophoblast, Epithelial-Mesenchymal Transition. Cells, 2025. 14(17).

22. Villarroel, F., et al., Exposure to fine particulate matter 2.5 from wood combustion smoke causes vascular changes in placenta and reduce fetal size. Reproductive Toxicology, 2024. 127: p. 108610.

23. Ziemendorff, A.C., et al., The moderating and mediating role of the placenta in the association between prenatal exposure to air pollutants and birth weight: A twin study. Environ Res, 2025. 270: p. 120952.

24. Furukawa, S., N. Tsuji, and A. Sugiyama, Morphology and physiology of rat placenta for toxicological evaluation. Journal of Toxicologic Pathology, 2018. 32: p. 1–17.

25. Kaufmann, P., S. Black, and B. Huppertz, Endovascular Trophoblast Invasion: Implications for the Pathogenesis of Intrauterine Growth Retardation and Preeclampsia. Biology of Reproduction, 2003. 69(1): p. 1–7.

26. Furukawa, S., et al., Toxicological Pathology in the Rat Placenta. Journal of Toxicologic Pathology, 2011. 24(2): p. 95–111.

27. Stapleton, P.A., et al., Maternal engineered nanomaterial exposure and fetal microvascular function: does the Barker hypothesis apply? American Journal of Obstetrics and Gynecology, 2013. 209(3): p. 227.e1–227.e11.

28. Stapleton, P.A., et al., Microvascular and mitochondrial dysfunction in the female F1 generation after gestational TiO2 nanoparticle exposure. Nanotoxicology, 2015. 9(8): p. 941–951.

29. Fournier, S.B., et al., Effect of Gestational Age on Maternofetal Vascular Function Following Single Maternal Engineered Nanoparticle Exposure. Cardiovascular Toxicology, 2019. 19(4): p. 321–333.

30. Yi, J., et al., Whole-body nanoparticle aerosol inhalation exposures. J Vis Exp, 2013(75): p. e50263.

31. Nurkiewicz, T.R., et al., Nanoparticle inhalation augments particle-dependent systemic microvascular dysfunction. Part Fibre Toxicol, 2008. 5: p. 1.

32. Inc., S.-A., Titanium (IV) oxide. 2024: St. Louis, MO.

33. Wang, K., et al., The effect of PM(2.5) exposure on placenta and its associated metabolites: A birth cohort study. Ecotoxicol Environ Saf, 2025. 292: p. 117891.

34. Böhm-González, S.T., et al., Association between trimester-specific prenatal air pollution exposure and placental weight of twins. Placenta, 2024. 154: p. 207–215.

35. Behlen, J.C., et al., Gestational Exposure to Ultrafine Particles Reveals Sex- and Dose-Specific Changes in Offspring Birth Outcomes, Placental Morphology, and Gene Networks. Toxicological Sciences, 2021. 184(2): p. 204–213.

36. Jones, M.L., et al., Antioxidant Defenses in the Rat Placenta in Late Gestation: Increased Labyrinthine Expression of Superoxide Dismutases, Glutathione Peroxidase 3, and Uncoupling Protein 21. Biology of Reproduction, 2010. 83(2): p. 254–260.

37. Tao, S., et al., Maternal exposure to ambient PM2.5 causes fetal growth restriction via the inhibition of spiral artery remodeling in mice. Ecotoxicology and Environmental Safety, 2022. 237: p. 113512.

38. Sandovici, I., et al., Placental adaptations to the maternal–fetal environment: implications for fetal growth and developmental programming. Reproductive biomedicine online, 2012. 25(1): p. 68–89.

39. Veras, M.M., et al., Particulate Urban Air Pollution Affects the Functional Morphology of Mouse Placenta1. Biology of Reproduction, 2008. 79(3): p. 578–584.

40. Hayashi, M., et al., Up-regulation of c-met protooncogene product expression through hypoxia-inducible factor-1α is involved in trophoblast invasion under low-oxygen tension. Endocrinology, 2005. 146(11): p. 4682–4689.

41. Graham, C.H., et al., Adriana and Luisa Castellucci Award Lecture 1999: Role of Oxygen in the Regulation of Trophoblast Gene Expression and Invasion. Placenta, 2000. 21(5): p. 443–450.

42. Griffith, J.A., et al., Maternal nano-titanium dioxide inhalation alters fetoplacental outcomes in a sexually dimorphic manner. Front Toxicol, 2023. 5: p. 1096173.

43. Cary, C.M., et al., Single pulmonary nanopolystyrene exposure in late-stage pregnancy dysregulates maternal and fetal cardiovascular function. Toxicol Sci, 2024. 199(1): p. 149–159.

44. Shukla, V. and M.J. Soares, Modeling Trophoblast Cell-Guided Uterine Spiral Artery Transformation in the Rat. Int J Mol Sci, 2022. 23(6).

45. Oberdörster, G., E. Oberdörster, and J. Oberdörster, Nanotoxicology: An Emerging Discipline Evolving from Studies of Ultrafine Particles. Environmental Health Perspectives, 2005. 113(7): p. 823–839.

