## Supplemental Figure 1 for "Gestational inhalation of nanoparticles disrupts placental zone structure and induces vascular placentation in rats"

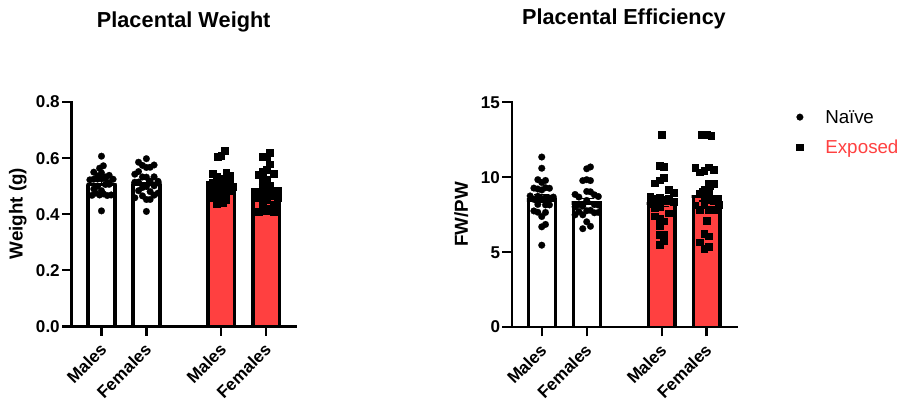

**Supplemental Figure 1.** Effects of nano-TiO_2_ inhalation on placental morphometric parameters. Placental weight (PW) (A) and placental efficiency (B) were measured on GD 20. Placental efficiency was calculated by dividing average fetal weight (FW) by average PW. Data are mean ± SEM, n=27-28 litters/treatment group. Analyzed by two-way ANOVA.
